# Characterization of nonlinear receptive fields of visual neurons by convolutional neural network

**DOI:** 10.1101/348060

**Authors:** Jumpei Ukita, Takashi Yoshida, Kenichi Ohki

## Abstract

A comprehensive understanding of the stimulus-response properties of individual neurons is necessary to crack the neural code of sensory cortices. However, a barrier to achieving this goal is the difficulty of analyzing the nonlinearity of neuronal responses. In computer vision, artificial neural networks, especially convolutional neural networks (CNNs), have demonstrated state-of-the-art performance in image recognition by capturing the higher-order statistics of natural images. Here, we incorporated CNN for encoding models of neurons in the visual cortex to develop a new method of nonlinear response characterization, especially nonlinear estimation of receptive fields (RFs), without assumptions regarding the type of nonlinearity. Briefly, after training CNN to predict the visual responses of neurons to natural images, we synthesized the RF image such that the image would predictively evoke a maximum response (“maximization-of-activation” method). We first demonstrated the proof-of-principle using a dataset of simulated cells with various types of nonlinearity, revealing that CNN could be used to estimate the nonlinear RF of simulated cells. In particular, we could visualize various types of nonlinearity underlying the responses, such as shift-invariant RFs or rotation-invariant RFs. These results suggest that the method may be applicable to neurons with complex nonlinearities, such as rotation-invariant neurons in higher visual areas. Next, we applied the method to a dataset of neurons in the mouse primary visual cortex (V1) whose responses to natural images were recorded via two-photon Ca^2+^ imaging. We could visualize shift-invariant RFs with Gabor-like shapes for some V1 neurons. By quantifying the degree of shift-invariance, each V1 neuron was classified as either a shift-variant (simple) cell or shift-invariant (complex-like) cell, and these two types of neurons were not clustered in cortical space. These results suggest that the novel CNN encoding model is useful in nonlinear response analyses of visual neurons and potentially of any sensory neurons.

## Author summary

A goal of sensory neuroscience is to comprehensively understand the stimulus-response properties of neuronal populations. However, a barrier to achieving this goal is the difficulty of analyzing the nonlinearity of neuronal responses, and existing methods for nonlinear response analyses are often designed to address specific types of nonlinearity of responses. In this study, we present a novel assumption-free method for nonlinear characterization of visual responses, especially nonlinear estimation of receptive fields (RFs), using a convolutional neural network (CNN), which has achieved state-of-the-art performance in computer vision. The proposed method was validated as follows. First, when trained to predict neuronal responses to natural images, the model yielded the best prediction accuracy among several machine-learning-based encoding models. Second, nonlinear RFs were successfully visualized from the trained CNN. Third, the shift-invariance of the responses, a well-known nonlinear property in V1 complex cells, was quantified from the visualized RFs. These results support the efficacy of a CNN encoding model for nonlinear response analyses that does not require explicit assumptions regarding the nonlinearity of neuronal responses. This study will contribute to the elucidation of nonlinear computations performed in neurons in the visual cortex and possibly any sensory cortex.

## Introduction

A goal of sensory neuroscience is to comprehensively understand the stimulus-response properties of neuronal populations. In the visual cortex, such properties were first characterized by Hubel and Wiesel, who discovered the orientation and direction selectivity of simple cells in the primary visual cortex (V1) using simple bar stimuli [1]. Later studies revealed that the responses of many visual neurons, including even simple cells [2–5], display nonlinearity, such as shift-invariance in V1 complex cells [6]; size, position, and rotation-invariance in inferotemporal cortex [7–9]; and viewpoint-invariance in a face patch [10]. Nevertheless, nonlinear response analyses of visual neurons have been limited thus far, and existing analysis methods are often designed to address specific types of nonlinearity underlying the neuronal responses. For example, the spike-triggered average [11] assumes linearity; moreover, the second-order Wiener kernel [12] and spike-triggered covariance [13–15] address second-order nonlinearity at most. In this study, we aim to analyze visual neuronal responses using an encoding model that does not assume the type of nonlinearity.

An encoding model that is useful for nonlinear response analyses of visual neurons must capture the nonlinear stimulus-response relationships of neurons. Thus, the model should be able to predict neuronal responses to stimulus images with high accuracy [16] even if the responses are nonlinear. In addition, the features that the encoding model represents should be visualized at least in part so that we can understand the neural computations underlying the responses. Artificial neural networks are promising candidates that may meet these criteria. Neural networks are mathematically universal approximators in that even one-hidden-layer neural network with many hidden units can approximate any smooth function [17]. In computer vision, neural networks trained with large-scale datasets have yielded state-of-the-art and sometimes human-level performance in digit classification [18], image classification [19], and image generation [20], demonstrating that neural networks, especially convolutional neural networks (CNNs) [21,22], capture the higher-order statistics of natural images through hierarchical information processing. In addition, recent studies in computer vision have provided techniques to extract and visualize the features learned in neural networks [23–26].

Several previous studies have used artificial neural networks as encoding models of visual neurons. These studies showed that artificial neural networks are highly capable of predicting neuronal responses with respect to low-dimensional stimuli such as bars and textures [27,28] or to complex stimuli such as natural stimuli [29–35]. Furthermore, receptive fields (RFs) were visualized by the principal components of the network weights between the input and hidden layer [29], by linearization [31], and by inversion of the network to evoke at most 80% of maximum responses [32]. However, these indirect RFs are not guaranteed to evoke the highest response of the target neuron.

In this study, we first investigated whether nonlinear RFs could be directly estimated by CNN encoding models (Fig 1) using a dataset of simulated cells with various types of nonlinearities. We confirmed that CNN yielded the best accuracy among several encoding models in predicting visual responses to natural images. Moreover, by synthesizing the image such that it would predictively evoke a maximum response (“maximization-of-activation” method), nonlinear RFs could be accurately estimated. Specifically, by repeatedly estimating RFs for each cell, we could visualize various types of nonlinearity underlying the responses without any explicit assumptions, suggesting that this method may be applicable to neurons with complex nonlinearities, such as rotation-invariant neurons in higher visual areas. Next, we applied the same procedures to a dataset of mouse V1 neurons, showing that CNN again yielded the best prediction accuracy among several encoding models and that shift-invariant RFs with Gabor-like shapes could be estimated for some cells from the CNNs. Furthermore, by quantifying the degree of shift-invariance of each cell using the estimated RFs, we classified V1 neurons as shift-variant (simple) cells and shift-invariant (complex-like) cells. Finally, these cells were not spatially clustered in cortical space. These results verify that nonlinear RFs of visual neurons can be characterized using CNN encoding models.

**Fig 1.**
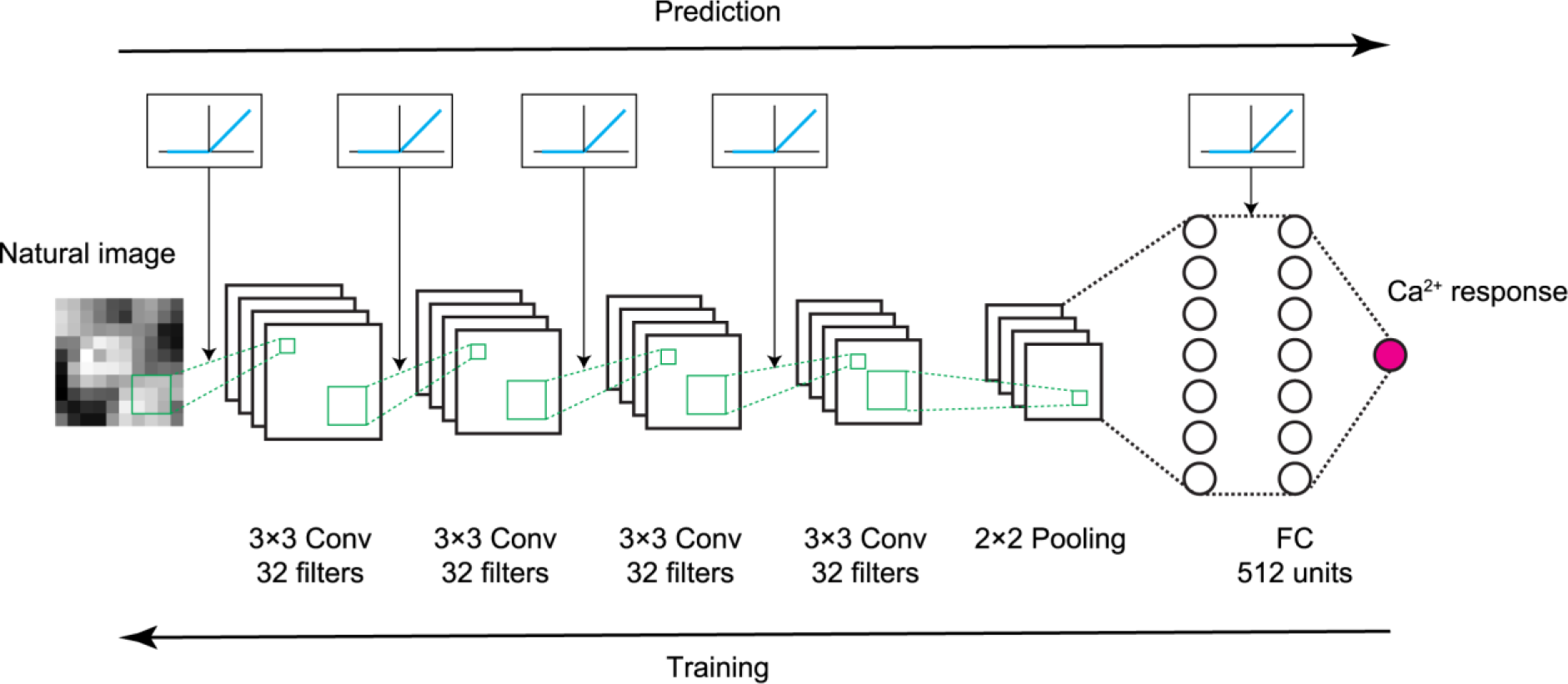
Scheme of CNN encoding model. The Ca^2+^ response to a natural image was predicted by convolutional neural network (CNN) consisting of 4 successive convolutional layers, one pooling layer, one fully connected layer, and the output layer (magenta circle). See Materials and Methods for details. Briefly, a convolutional layer calculates a 3×3 convolution of the previous layer followed by a rectified linear (ReLU) transformation. The pooling layer calculates max-pooling of 2×2 regions in the previous layer. The fully connected layer calculates the weighted sum of the previous layer followed by a ReLU transformation. The output layer calculates the weighted sum of the previous layer followed by a sigmoidal transformation. During training, parameters were updated by backpropagation to reduce the mean squared error between the predicted responses and actual responses.

## Results

### Nonlinear RFs could be estimated by CNN encoding models for simulated cells with various types of nonlinearities

We generated a dataset comprising the stimulus natural images (2200 images) and the corresponding responses of simulated cells. To investigate the ability of CNN to handle various types of nonlinearities, we incorporated various basic nonlinearities for the data generation, including rectification, shift-invariance, and in-plane rotation-invariance, which were found in V1 simple cells [2], V1 complex cells [6], and inferotemporal cortex [9], respectively. We generated the responses of simple cells (N = 30), complex cells (N = 70), and rotation-invariant cells (N = 10) using the linear-nonlinear model [2], energy model [36,37], and rotation-invariant model, respectively (Figs 2A, 2B, and 3A; see Materials and Methods for details). The responses were generated using one Gabor-shaped filter for a simple cell, two phase-shifted Gabor-shaped filters for a complex cell, and 36 rotated Gabor-shaped filters for a rotation-invariant cell. We also added some noise sampled from a Gaussian distribution such that the trial-to-trial variability of simulated data was similar to that of real data.

**Fig 2.**
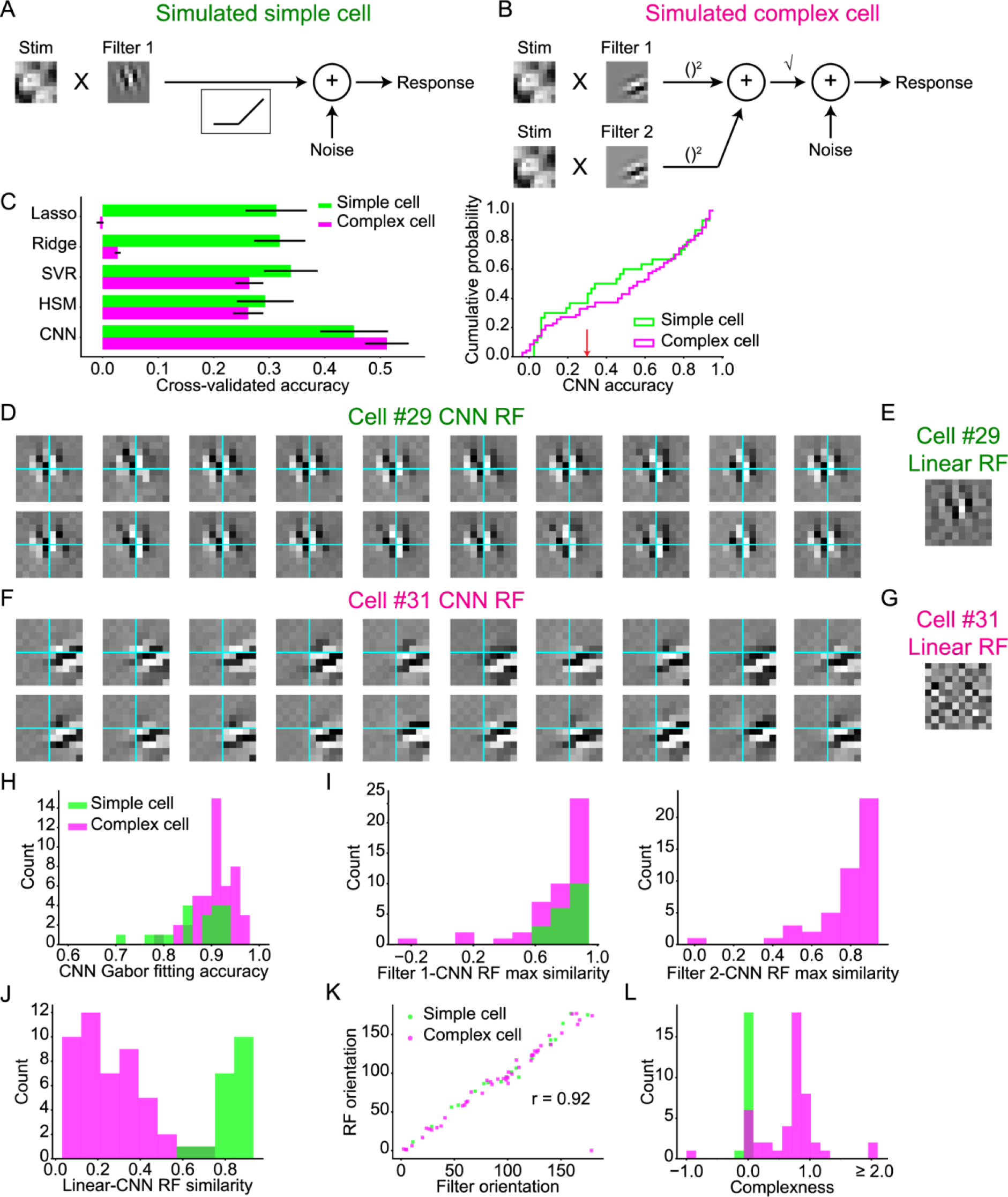
Nonlinear RFs could be estimated by CNN encoding models for simulated simple cells and complex cells. (A) Scheme of response generation for simulated simple cells. The response to a stimulus was defined as the rectified dot product between the stimulus image and a Gabor-shaped filter, followed by an additive Gaussian noise. The Gabor-shaped filter of simulated simple cell #29 is displayed in this panel. (B) Scheme of response generation for simulated complex cells. The response to a stimulus was defined as the square root of the squared sum of the output of two subunits, followed by an additive Gaussian noise. Each subunit, which had a Gabor-shaped filter with a shifted phase, calculated the dot product between the stimulus image and the filter (See Materials and Methods for details). The Gabor-shaped filters of simulated complex cell #31 are displayed in this panel. (C) Left: comparison of the response prediction accuracies among the following encoding models: the L1-regularized linear regression model (Lasso), L2-regularized linear regression model (Ridge), support vector regression model (SVR), hierarchical structural model (HSM), and CNN. Data are presented as the mean ± s.e.m. (N = 30 simulated simple cells and N = 70 simulated complex cells). Right: cumulative distribution of CNN prediction accuracy. Simulated cells with a CNN prediction accuracy ≤ 0.3 (indicated as the red arrow) were removed from the following receptive field (RF) analysis. (D, F) Results of iterative CNN RF estimations for simulated simple cell #29 (D) and complex cell #31 (F). Only 20 of the 100 generated RF images are shown in these panels. Grids are depicted in cyan. Although the simulated simple cell #29 had RFs in nearly identical positions, the simulate complex cell #31 had RFs in shifted positions. (E, G) Linearly estimated RFs (linear RFs) of simulated simple cell #29 (E) and complex cell #31 (G), using a regularized pseudoinverse method. (H) Gabor-fitting accuracy of CNN RFs. Accuracy was defined as the Pearson correlation coefficient between the CNN RF and fitted Gabor kernel. (I) Maximum similarity between each generator filter and 100 CNN RFs. (J) Similarity between linear RFs and CNN RFs. Similarity was defined as the normalized pixelwise dot product between the linear RF and CNN RF. (K) Relationship of the Gabor orientations between generator filters and CNN RFs. (L) Distribution of complexness. Only cells with a CNN prediction accuracy > 0.3 were analyzed in H-L (N = 19 simple cells and N = 47 complex cells).

**Fig 3.**
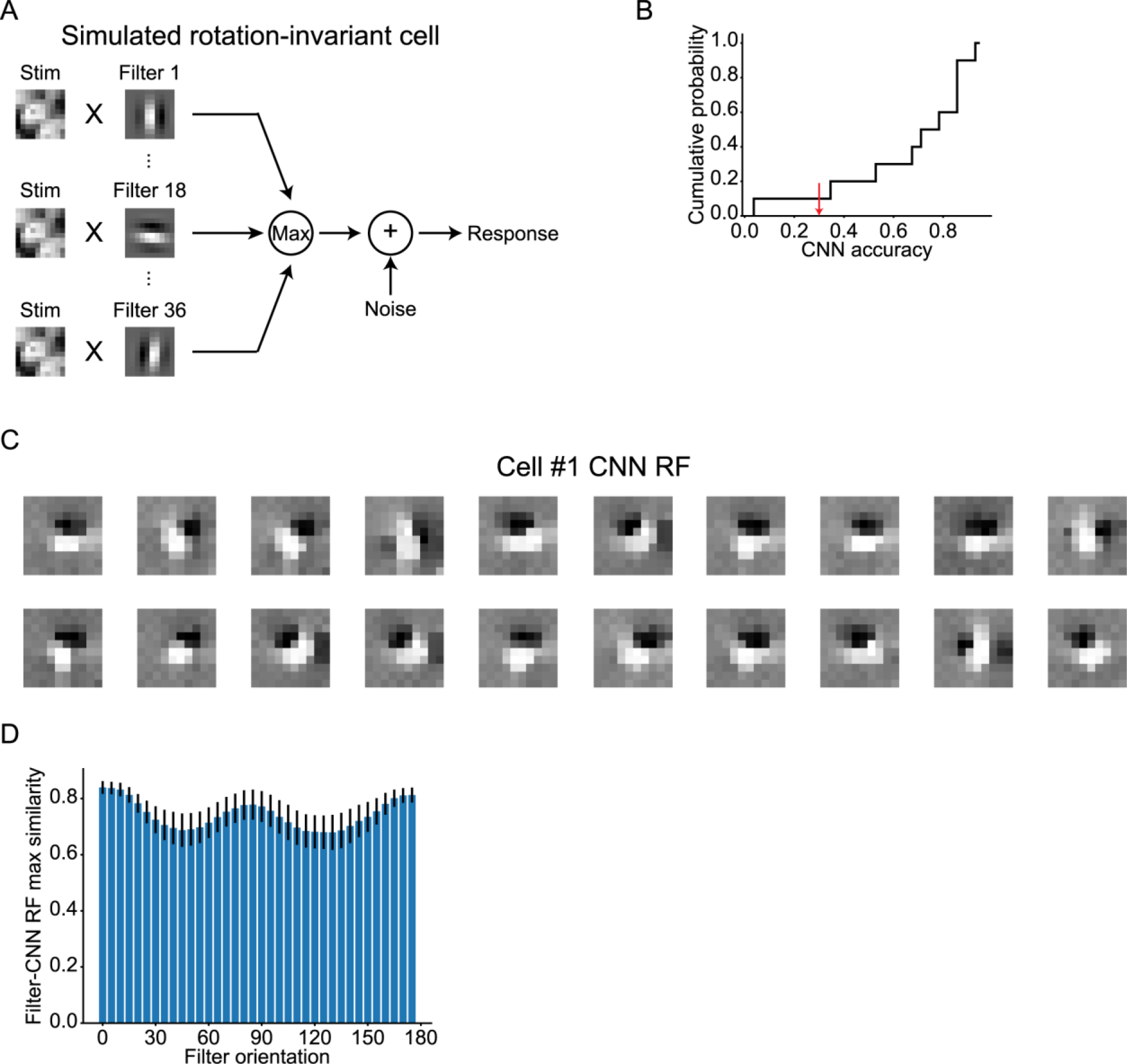
Nonlinear RFs could be estimated by CNN encoding models for simulated rotation-invariant cells. (A) Scheme of response generation for simulated rotation-invariant cells. The response to a stimulus was defined as the maximum of the output of 36 subunits followed by an additive Gaussian noise. Each subunit, which had a Gabor-shaped filter with different orientations, calculated the dot product between the stimulus image and the filter (See Materials and Methods for details). The filters of simulated cell #1 are displayed in this panel. (B) Cumulative distribution of CNN prediction accuracy (N = 10 cells). Simulated cells with a CNN prediction accuracy ≤ 0.3 (indicated as the red arrow) were removed from the following RF analysis. (C) Results of iterative CNN RF estimations for simulated cell #1. Only 20 of the 1000 generated RF images are shown in this panel. RF images had Gabor-like shapes but their orientations were different in different iterations. (D) Maximum similarity between each generator filter and 1000 CNN RFs. Only cells with a CNN prediction accuracy > 0.3 were analyzed (N = 9 cells).

We first used a dataset of simulated simple cells and complex cells and trained the CNN for each cell to predict responses with respect to the natural images (Fig 1). For comparison, we also constructed the following types of encoding models: an L1-regularized linear regression model (Lasso), L2-regularized linear regression model (Ridge), support vector regression model (SVR) with a radius basis function kernel, and hierarchical structural model (HSM) [31]. The prediction accuracy, defined as the Pearson correlation coefficient between the predicted responses and actual responses in a 5-fold cross-validation manner, of CNN was high and better than that of other models for both simple cells and complex cells (Fig 2C), ensuring that the stimulus-response relationships of these cells were successfully captured by CNN.

Next, we visualized the RF of each cell using the maximization-of-activation approach (see Materials and Methods) [23,24] where the RF was regarded as the image that evoked the highest activation of the output layer of the trained CNN. We performed this RF estimation 100 times independently for each cell, utilizing the empirical fact that an independent iteration of RF estimation processes creates different RF images by finding different maxima [23]. Fig 2D and 2F show 20 out of the 100 RF images estimated by the trained CNN (CNN RF images) for a representative simple cell and complex cell, respectively. The predicted responses with respect to these RF images were all > 99% of the maximum response in the actual data of each cell, ensuring that the activations of the CNN output layers were indeed maximized. All visualized RF images had clearly segregated ON and OFF subregions, and the structure was close to the Gabor-shaped filters used in the response generations (Fig 2D vs. Fig 2A and Fig 2F vs. Fig 2B). Furthermore, when RF images were compared within a cell, RF images of cell #29 had ON and OFF subregions in nearly identical positions, while some RF images of cell #31 were shifted in relation to one another. These observations are consistent with the assumption that cell #29 is a simple cell and cell #31 is a complex cell.

For complex cells, we expect that RF estimation using linear methods would fail to generate an image with clearly segregated ON and OFF subregions, whereas nonlinear RF estimation would not [14]. Thus, the similarity between a linearly estimated RF image (linear RF) and a nonlinearly estimated RF image is expected to be low for complex cells. We performed linear RF estimations following a previous study [38]. Although the linear RF image and CNN RF image were similar for cell #29 (Fig 2E), the linear RF image for cell #31 was ambiguous, lacked clear subregions, and was in sharp contrast to the CNN RF image (Fig 2G). These results are again consistent with the assumption that cell #29 is a simple cell and cell #31 is a complex cell.

Next, we comprehensively analyzed the RFs of populations of simulated simple cells and complex cells. Cells with a CNN prediction accuracy ≤ 0.3 were omitted from the analyses (Fig 2C). First, the similarity between a linear RF image and CNN RF image, measured as the maximum normalized pixelwise dot product between a linear RF image and 100 CNN RF images, was distinctly different between simple cells and complex cells (Fig 2J), reflecting different degrees of nonlinearity. Second, the accuracy of Gabor-kernel fitting of the CNN RF image, measured as the pixelwise Pearson correlation coefficient between a CNN RF image and the fitted Gabor kernel, was high among all analyzed cells (Fig 2H), confirming that the estimated RFs had a shape similar to a Gabor kernel. Third, the maximum similarity between each filter used in the response generation and 100 CNN RF images were high for both simple cells and complex cells (Fig 2I). Fourth, the orientations of the CNN RF images, estimated by fitting them to Gabor kernels, were nearly identical to the orientations of the filters of the response generators (circular correlation coefficient [39] = 0.92; Fig 2K). These results suggest that the RFs estimated by the CNN encoding models had similar structure to the ground truth and that the shift-invariant property of complex cells was successfully visualized from iterative RF estimations.

We also performed similar analyses using a dataset of simulated rotation-invariant cells. When trained to predict the responses with respect to the natural images, CNNs again yielded high prediction accuracy (Fig 3B). Next, we estimated RFs using the maximization-of-activation approach independently 1000 times for each cell. The predicted responses with respect to these RF images were all > 99% of the maximum response in the actual data of each cell, ensuring that the activations of CNN output layers were indeed maximized. As shown in Fig 3C, the visualized RF images of cell #1 had Gabor shapes close to the filters used in the response generation (Fig 3A). In addition, some RF images were rotated in relation to one another, consistent with the rotation-invariant response property of this cell. Finally, we quantitively compared the RFs (1000 RF images for each cell) and the filters of the response generator (36 filters for each cell). For each filter, the maximum similarity with 1000 CNN RF images was high (Fig 3D), suggesting that the estimated RFs had various orientations and similar structure to the ground truth. Thus, using the proposed RF estimation approach, RFs were successfully estimated by the CNN encoding models, and various types of nonlinearity could be visualized from multiple RFs without assumptions, although the hyperparameters and layer structures of CNNs were unchanged across cells.

### CNN yielded the best accuracy for prediction of the visual response of V1 neurons

Next, we used a dataset comprising the stimulus natural images (200-2200 images) and corresponding real neuronal responses (N = 2465 neurons, 4 planes), which were recorded using two-photon Ca^2+^ imaging from mouse V1 neurons. To investigate whether CNN was able to capture the stimulus-response relationships of V1 neurons, we trained the CNN for each neuron to predict the neuronal responses to the natural images (Fig 1). The prediction accuracy was again measured by the Pearson correlation coefficient between the predicted responses and actual responses of the held-out test data in a 5-fold cross-validation manner (N = 2455 neurons that were not used for the hyperparameter optimizations; see Materials and Methods). Comparison of the prediction accuracies among several types of encoding models revealed that CNN outperformed other models (Fig 4A), and the prediction of the CNNs were accurate (Fig 4B and 4C). These results show that the stimulus-response relationships of V1 neurons were successfully captured by CNN, demonstrating the efficacy of using CNN for further RF analyses of V1 neurons.

**Fig 4.**
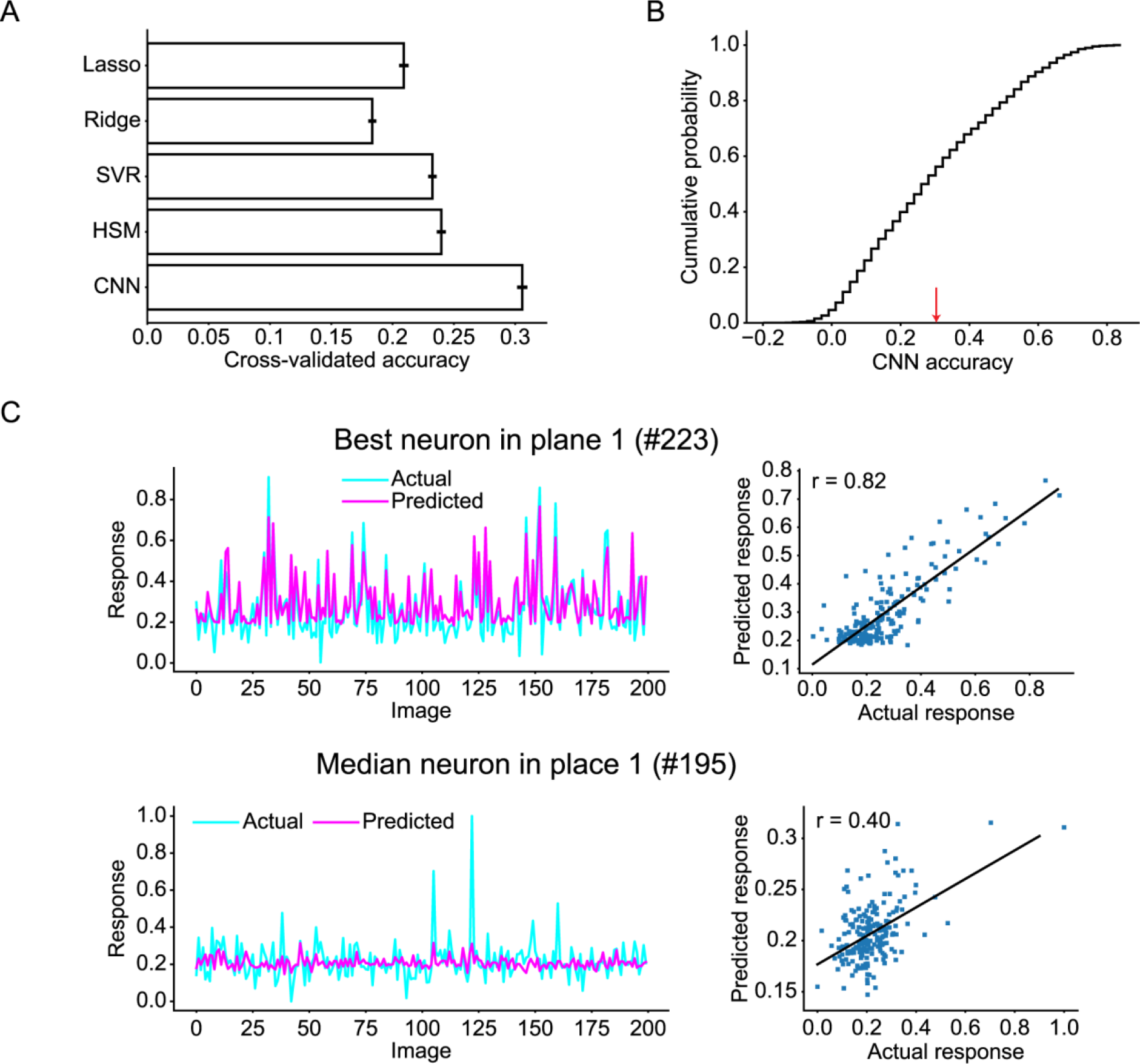
Prediction accuracy of the CNN for V1 neurons. (A) Comparison of the response prediction accuracies among various encoding models: the L1-regularized linear regression model (Lasso), L2-regularized linear regression model (Ridge), SVR, HSM, and CNN. Data are presented as the mean ± s.e.m. (N = 2455 neurons). (B) Cumulative distribution of CNN prediction accuracy. Neurons with a CNN prediction accuracy ≤ 0.3 (indicated as the red arrow) were removed from the following RF analysis. (C) Distributions of actual responses and predicted responses of the neuron with the best prediction accuracy in a plane (top) and the neuron with the median prediction accuracy in a plane (bottom). Each dot in the right panel indicates data for each stimulus image. Solid lines in the right panels are the linear least-squares fit lines. Only data for 200 images are shown.

### Estimation of nonlinear RFs of V1 neurons from CNN encoding models

Next, we visualized the RF of each neuron by the maximization-of-activation approach (see Materials and Methods) [23,24]. Neurons with a CNN prediction accuracy ≤ 0.3 were omitted from this analysis (Fig 4B). The resultant RF images for two representative neurons are shown in Fig 5B. Both RF images have clearly segregated ON and OFF subregions and were well fitted with two-dimensional Gabor kernels (Fig 5C), consistent with known characteristics of simple cells and complex cells in V1 [14,40]. The accuracy of Gabor-kernel fitting, measured as the pixelwise Pearson correlation coefficient between the RF image and fitted Gabor kernel, was high among all analyzed neurons (median r = 0.77; Fig 5E), suggesting that the RF images generated from the trained CNNs (CNN RF images) successfully captured the Gabor-like structure of RFs observed in V1. We also performed linear RF estimations following a previous study [38]. Although the linear RF image and CNN RF image were similar for neuron #639, the linear RF image for neuron #646 was ambiguous, lacked clear subregions, and was in sharp contrast to the CNN RF image (Fig 5A and 5B), suggesting that neuron #639 would be linear and neuron #646 would be nonlinear. Supporting this idea, further analysis (see below) revealed that neuron #639 was a shift-variant (simple) cell, and neuron #646 was a shift-invariant (complex-like) cell. The similarity between a linear RF image and a CNN RF image, measured as the normalized pixelwise dot product between these two images, varied among all analyzed neurons (Fig 5D), reflecting the distributed nonlinearity of V1 neurons.

**Fig 5.**
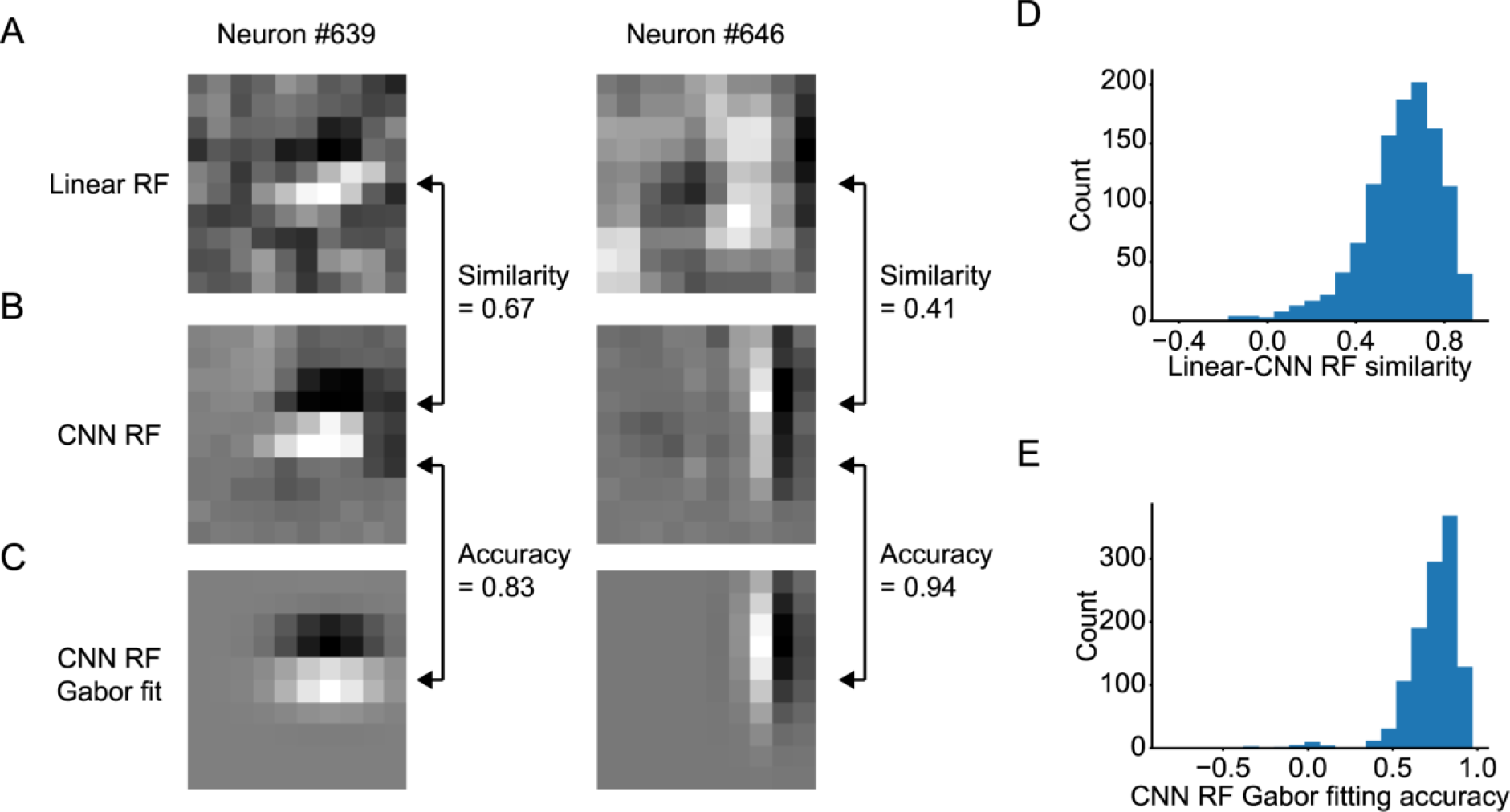
Estimating RFs of V1 neurons from trained CNNs. (A) Linearly estimated RFs (linear RFs) of two representative neurons (#639 and #646), using a regularized pseudoinverse method. (B) RFs estimated from the trained CNNs (CNN RFs) of the two representative neurons. (C) Gabor kernels fitted to CNN RFs of the two representative neurons. (D) Similarity between linear RFs and CNN RFs. Similarity was defined as the normalized pixelwise dot product between the linear RF and the CNN RF. (E) Gabor fitting accuracy of CNN RFs. Accuracy was defined as the Pearson correlation coefficient between the CNN RF and the fitted Gabor kernel. Only neurons with a CNN prediction accuracy > 0.3 were analyzed in D and E (N = 1160 neurons).

### Estimated RFs of some V1 neurons were shift-invariant

We then performed 100 independent CNN RF estimations for each V1 neuron to characterize the nonlinearity of RFs. We especially focused on the shift-invariance, the most well-studied nonlinearity in V1 complex cells [6]. Fig 6 shows 20 of the 100 CNN RF images for two representative neurons. The predicted responses with respect to these RF images were all > 99% of the maximum response in the actual data of each neuron, ensuring that the activations of the CNN output layers were indeed maximized. Importantly, RF images of neuron #639 had ON and OFF subregions in nearly identical positions (Fig 6A). In contrast, some RF images of neuron #646 were horizontally shifted in relation to one another (Fig 6B), suggesting that neuron #646 is shift-invariant and could be a complex cell.

**Fig 6.**
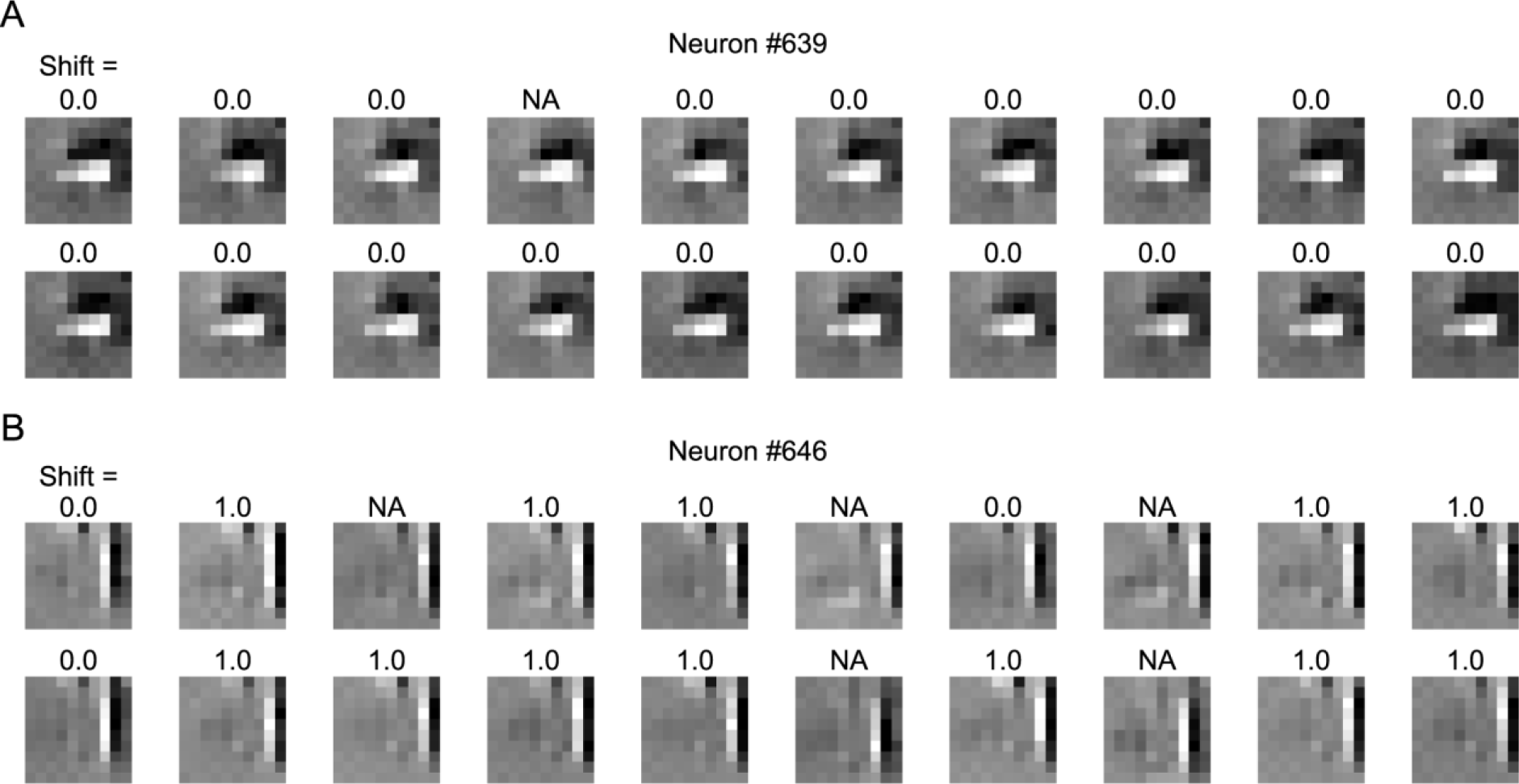
Examples of iterative CNN RF estimation for V1 neurons. Results of iterative CNN RF estimations for neuron #639 (A) and neuron #646 (B). Only 20 out of the 100 generated RF images are shown in this figure. The number above each RF image indicates the shift pixel distance between the RF image and the top left RF image. The shift distance between two images was calculated as the maximum distance of pixel shifts with which the zero-mean normalized cross correlation (ZNCC) > 0.95, projected orthogonally to the Gabor orientation. “NA” indicates that the ZNCC was not above 0.95 for any shift. While shift distances were zero or NA for RF images of neuron #639, some RF images of neuron #646 were shifted to another by one pixel.

### Characterization of shift invariance from iteratively estimated RF images

To quantitatively understand the shift-invariance, we then developed predictive models of visual responses for each simulated complex cell and V1 neuron, termed simple model and complex model, inspired by the stimulus-response properties of simple and complex cells. In the simple model, the response to a stimulus was predicted as the normalized dot product between the stimulus image and an RF image. The RF image that yielded the best prediction accuracy was chosen and used for all stimulus images (Fig 7A). In contrast, in the complex model, the response to each stimulus was predicted as the maximum of the normalized dot products between the stimulus image and several RF images (Fig 7B). Here, RF images used in these models were selected from 100 RF images as ones that were shifted to one another. If there was no shifted RF image, the complex model was identical to the simple model (see Materials and Methods). Fig 7 shows examples of predictions from the simple and complex models for V1 neuron #646. Although the response to one image (Stim 1) was predicted moderately well by both the simple model and complex model, the prediction for another image (Stim 2) by the simple model was far poorer than the prediction by the complex model. This difference is probably because the ON/OFF phase of the RF image used in the simple model (RF 4) did not match with that of Stim 2. On the other hand, the complex model had multiple RF images, and one RF image (RF 1) matched with Stim 2. These results suggest that the responses of this neuron are somewhat tolerant to phase shifts and that such complex cell-like properties were better captured by the complex model than by the simple model.

**Fig 7.**
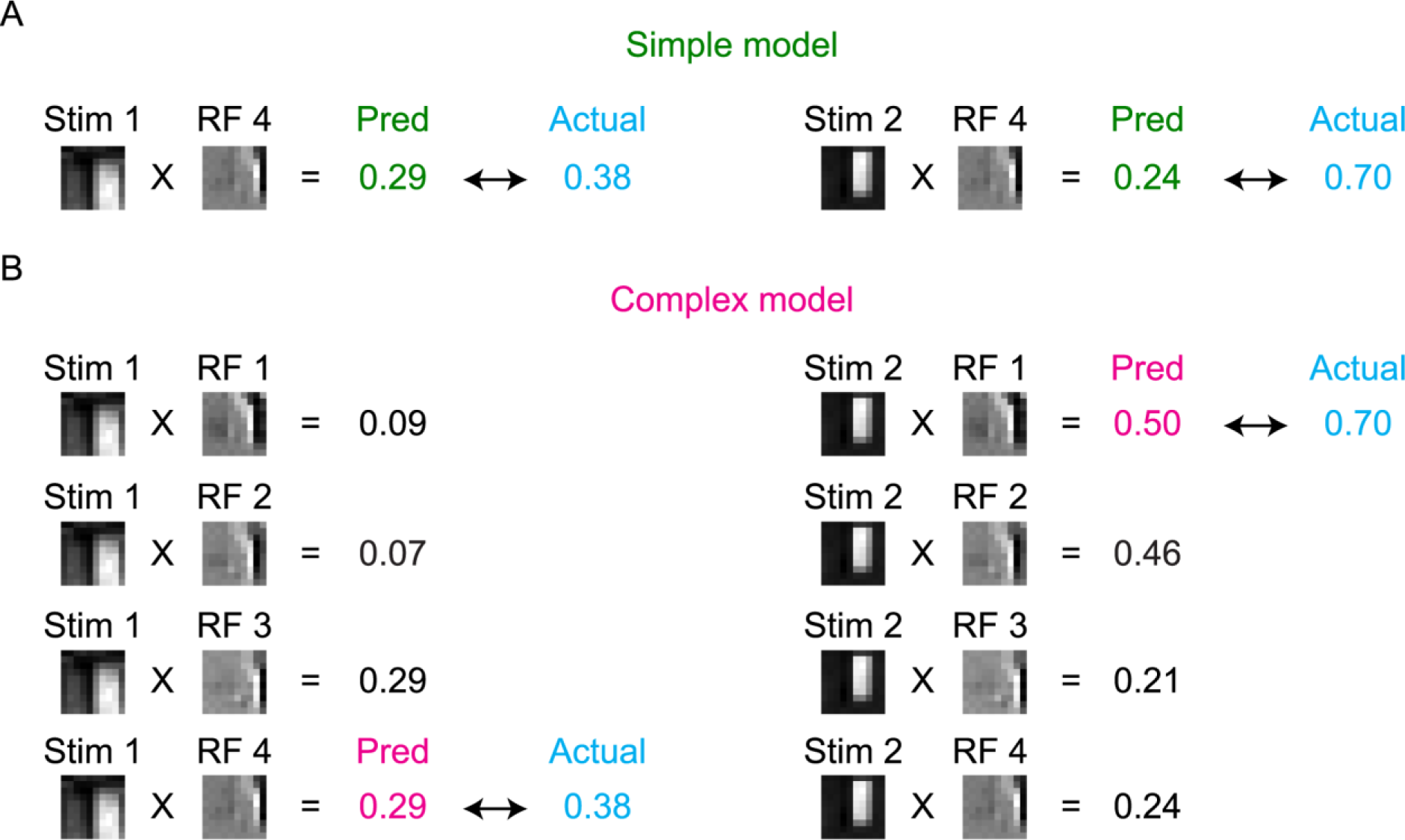
Schemes of the simple model and complex model. Schemes of the simple model and complex model are illustrated using RFs and actual responses of neuron #646. (A) The simple model is a linear predictive model, which predicts the neuronal response as the normalized dot product between the stimulus image and one RF image (RF 4). (B) The complex model predicts the neuronal response as the maximum of the normalized dot products of the stimulus image and several RF images (RF 1-4). Note that the complex model predicted the neuronal response to Stim 2 better than the simple model for this neuron.

We then measured the prediction accuracy of each model for all stimulus images by the Pearson correlation coefficient between the predicted responses and actual responses. As expected, the accuracy of the complex model was better than that of the simple model for this neuron #646 (Fig 8A and 8B), reflecting its shift-invariant property (Figs 5, 6 and 7).

**Fig 8.**
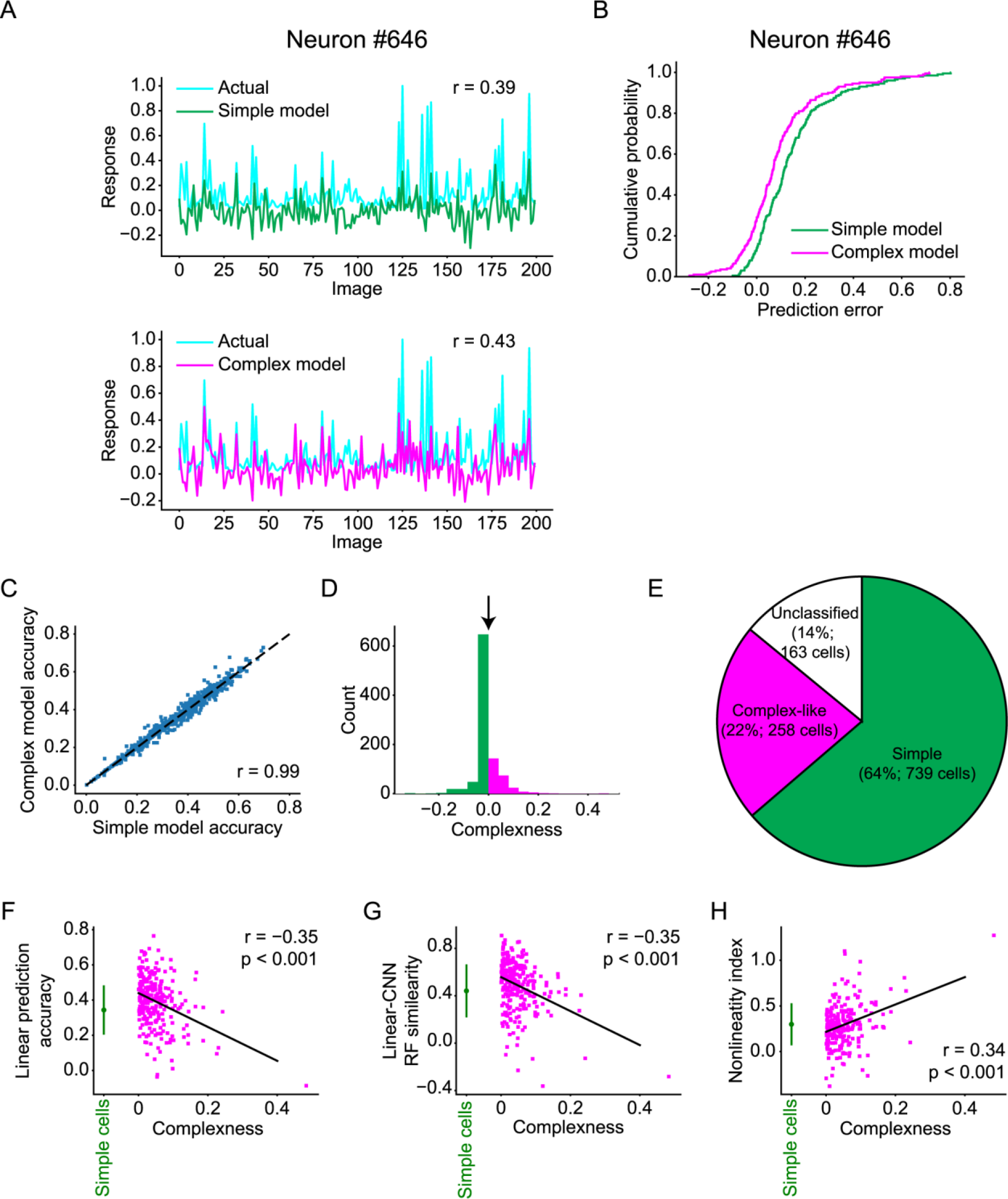
Simple cells and complex-like cells. (A) Distributions of the actual responses (cyan lines) and responses predicted by the simple model (green line in the top panel) and the complex model (magenta line in the bottom panel) for neuron #646. (B) Cumulative distributions of prediction errors of the simple model (green) and the complex model (magenta) for neuron #646. Prediction error was defined as the difference between the predicted response and actual response. (C) Relationship of accuracies between the simple model and complex model (N = 997 neurons). Neurons with the Gabor fitting accuracy ≤ 0.6, accuracy of the simple model < 0, or accuracy of the complex model < 0 were omitted from this analysis. (D) Distribution of complexness. Simple cells (green) and complex-like cells (magenta) were classified with threshold = 0 (black arrow). (E) Proportion of classified cells, simple cells, and complex-like cells among neurons with the CNN response prediction accuracy > 0.3. Classified cells were neurons with the Gabor fitting accuracy > 0.6, the response prediction accuracy of the simple model > 0, and the response prediction accuracy of the complex model > 0. Simple cells were neurons with complexness ≤ 0. Complex-like cells were neurons with complexness > 0. (F-H) Relationships between complexness and linear (Lasso) prediction accuracy (F), similarity between linear RFs and CNN RFs (G), and the nonlinearity index (H). Data of simple cells are presented as the mean ± s.d. (N = 739 neurons, green). Solid lines are the linear least-squares fit lines for complex-like cells. Both linear prediction accuracy and RF similarity of complex-like cells (magenta) negatively correlated with complexness (r = −0.35, p < 0.001, N = 258 neurons: F and r = −0.29, p < 0.001, N = 258 neurons: G), while the nonlinearity index of complex-like cells positively correlated with complexness (r = 0.34, p < 0.001, N = 258 neurons: H), suggesting that complexness defined here indeed reflected nonlinearity.

We compared the accuracy of the simple model and complex model for populations of V1 neurons (Fig 8C), simulated simple cells, and simulated complex cells. We defined the complexness index for each cell by

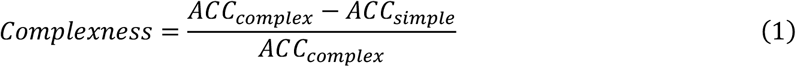

where *ACC*_*simpie*_ and *ACC*_*complex*_ are the response prediction accuracy of the simple model and complex model, respectively. Cells with a Gabor fitting accuracy (Figs 2H and 5E) ≤ 0.6, *ACC*_*simpie*_ < 0, or *ACC*_*complex*_ < 0 were omitted from this analysis. Then, we defined simple cells as cells with complexness ≤ 0 and complex-like cells as cells with complexness > 0. The sensitivity (recall) of this classification for simulated data was 89% for simple cells and 85% for complex cells (Fig 2L), ensuring the validity of this classification. In addition, the ratio of complex-like cells (26%, 258/997 neurons; Fig 8D and 8E) among V1 neurons was consistent with that in a previous study [41].

We also compared complexness with other indices of linearity and nonlinearity using a dataset of V1 neurons. First, linear prediction accuracy, measured as the prediction accuracy of the L1-regularized linear regression model (Lasso), significantly anti-correlated with complexness for complex-like cells (Fig 8F) (r = −0.35, p < 0.001, N = 258; Student’s t-test), suggesting that the linear regression models could not accurately predict the responses of neurons with high complexness. Similarity between linear RF images and CNN RF images also anti-correlated significantly with complexness (Fig 8G) (r = −0.35, p < 0.001, N = 258; Student’s t-test), suggesting that linear RFs could not accurately capture the RFs of neurons with high complexness. Furthermore, the nonlinearity index ((CNN prediction accuracy - Lasso prediction accuracy) / CNN prediction accuracy; see Materials and Methods) significantly correlated with complexness (Fig 8H) (r = 0.34, p < 0.001, N = 258, Student’s t-test), suggesting that the nonlinearity of V1 neurons was at least in part introduced by the nonlinearity of complex-like cells.

### Simple cells and complex-like cells were not spatially clustered in V1

Finally, we tested whether simple cells and complex-like cells were spatially organized in the cortical space. We first investigated the spatial structure of complexness by comparing the difference in complexness with the cortical distance between all neuron pairs (N = 129451 neuron pairs). We found no correlation between complexness and cortical distance (r = −0.01), suggesting no distinct spatial organization of complexness (Fig 9A left and B). We also calculated the cortical distances of all simple cell-simple cell pairs and complex-like cell-complex-like cell pairs. The cumulative distributions of these distances, normalized by the area, were both within the first and 99^th^ percentiles of the position-permuted simulations (1000 times for each plane; see Materials and Methods for the permutations), demonstrating no cluster organization of simple cells or complex-like cells (Fig 9 right and 9B).

**Fig 9.**
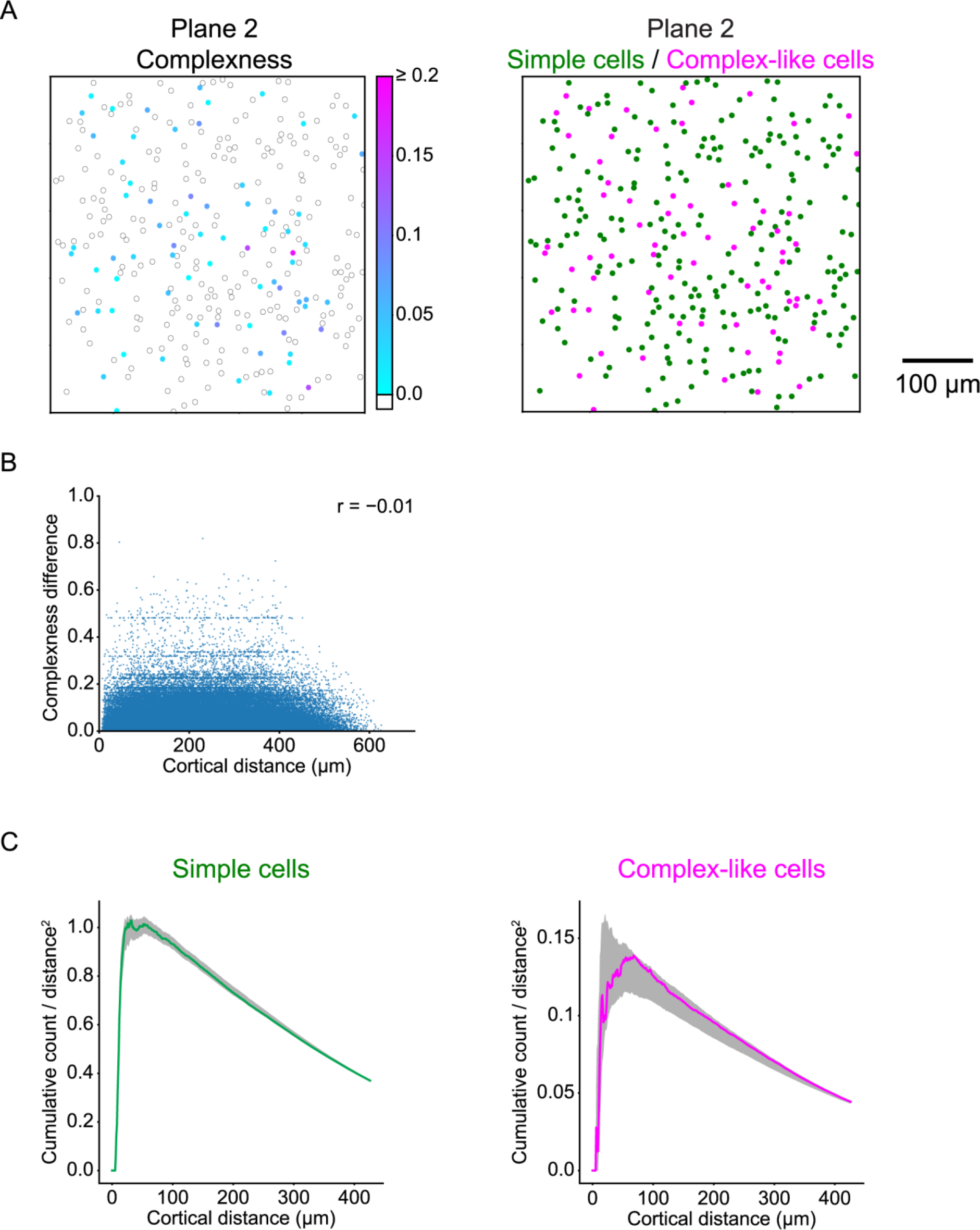
Spatial organizations of simple cells and complex-like cells. (A) Left: cortical distribution of complexness for the representative plane. Position of each neuron is represented as the circle annotated by the complexness (cyan to magenta for complex-like cells (complexness > 0) and white for simple cells (complexness ≤ 0)). Right: cortical distribution of simple cells (N = 238 neurons, green) and complex-like cells (N = 70 neurons, magenta) for the representative plane. (B) Relationship between cortical distances and differences of complexness for all simple cells and complex-like cells. (C) Cumulative distributions of the number of simple cell-simple cell pairs (left) or complex-like cell-complex-like cell pairs (right) as a function of the cortical distance, normalized by the area. Dark shadows indicate the range from the first to 99^th^ percentile of 1000 position-permuted simulations for each plane. The cumulative distributions were both within the first and 99^th^ percentiles of simulations, indicating no distinct spatial arrangements of simple cells or complex-like cells.

## Discussion

### Estimation of nonlinear RFs from CNN encoding models

We first revealed that the accuracy of CNN in predicting responses to natural images was high for both simulated cells and V1 neurons (Figs 2C, 3B, 4B). This finding is not surprising in light of the recent successes of artificial neural networks, especially CNN, in computer vision [18–20]. Such successes could be attributed to the ability of CNN to acquire sophisticated statistics of high-dimensional data [42]. Likewise, the high prediction accuracy of CNN shown in this study is possibly due to its ability to capture higher-order nonlinearity between stimulus images and responses. Notably, the prediction accuracy of CNN was high even though the hyperparameters and layer structures of CNNs were identical for all types of cells, suggesting that CNN might be used as a general-purpose encoding model of visual neurons.

Using simulated cells, we showed that nonlinear RFs could be accurately estimated by CNN encoding models by the maximization-of-activation approach. In particular, various types of response nonlinearity could be visualized, including RFs with different phases for complex cells (Figs 2D, 2F) and RFs with different orientations for rotation-invariant cells (Fig 3C). One advantage of this RF estimation method is that it does not require an explicit assumption regarding the nonlinearities of RFs, whereas most methods for nonlinear RF estimation in previous studies do. Second-order Wiener kernel [12] and spike-triggered covariance [13–15] are capable of estimating RFs with second-order nonlinearity at most, and Fourier-based methods [43,44] estimate RFs that are linearized in the Fourier domain. The second advantage is that our method can directly visualize the image that is predicted to evoke the highest response of the target cell, in contrast to previously proposed RF estimations from artificial neural networks [29,31,32]. As suggested in [45], the disadvantage of the maximization-of-activation approach is that it may produce unrealistic images even if the maximization of activation was successful because the candidate image space is extremely vast. To avoid this issue, we constrained the candidate image space to natural images by using L_p_-norm and total variance regularizations. Although the hyperparameters of regularizations were fixed across all analyzed cells, these regularizations worked well when considering the quality of the resultant RF images.

We then applied the RF estimation method to a dataset of V1 neurons and revealed that shift-invariant RFs could be estimated for complex-like cells from CNNs. Although direct quantification of the shift-invariant property of each cell from these RF images (e.g., by calculating the maximum shift distance orthogonal to the Gabor orientation) is indeed possible, it could lead to incorrect conclusions since the prediction accuracies of CNNs were imperfect (Figs 2C and 4B). For example, a CNN trained with low accuracy for a simple cell might not accurately implement the stimulus-response relationship of this cell and might accidentally generate some shifted RF images. Instead, the complexness was calculated as the difference in accuracies of the simple model and complex model (Figs 7 and 8) so that the complexness reflects the stimulus-response statistics of the data.

### Association between animal vision and deep learning

Although artificial neural networks and cortical neural networks have much in common [46], the former might not be an exact *in silico* implementation of the latter (e.g., the learning algorithms discussed in [47]). However, recent studies have suggested that the representations of CNNs and the activity of the visual cortex share hierarchical similarities [48–52]. These studies raise the possibility that the CNN encoding model could be applicable to neurons with complex nonlinearities, such as rotation-invariant neurons in the inferotemporal cortex [9]. Thus, the CNN encoding model and nonlinear RF characterization proposed in this paper will contribute to future studies of neural computations not only in V1 but also in higher visual areas.

## Materials and methods

### Acquisition of neural data

All experimental procedures were performed using C57BL/6 male mice (N = 3; Japan SLC, Hamamatsu, Shizuoka, Japan), which were approved by the Animal Care and Use Committee of Kyushu University and the University of Tokyo. Anesthesia was induced and maintained with isoflurane (5% for induction, 1.5% during surgery, and ~0.5% during imaging with a sedation of ~0.5 mg/kg chlorprothixene; Sigma-Aldrich, St Louis, MO, USA). After the skin was removed from the head, a custom-made metal head plate was attached to the skull with dental cement (Super Bond; Sun Medical, Moriyama, Shiga, Japan), and a craniotomy was made over V1 (center position: 0-1 mm anterior from lambda, +2.5-3 mm lateral from midline). Then, 0.8 mM Oregon green BAPTA-1 (OGB-1; Life Technologies, Grand Island, NY, USA), dissolved with 10% Pluronic (Life Technologies) and 25 μM sulforhodamine 101 (SR101; Sigma-Aldrich) was pressure-injected using Picospritzer III (Parker Hannifin, Cleveland, OH, USA) approximately 400 μm below the cortical surface. The craniotomy was sealed with a coverslip and dental cement.

Neuronal activity was recorded using two-photon microscopy (A1R MP; Nikon, Minato-ku, Tokyo, Japan) with a 25× objective lens (NA = 1.1; PlanApo, Nikon) and Ti:Sapphire mode-locked laser (Mai Tai DeepSee; Spectra Physics, Santa Clara, CA, USA). OGB-1 and SR101 were both excited at a wavelength of 920 nm, and their emissions were filtered at 525/50 nm and 629/56 nm, respectively. 507×507 μm or 338×338 μm images were obtained at 30 Hz using a resonant scanner with a 512 ×512-pixel resolution.

Visual stimuli were presented using PsychoPy [53] on a 32-inch LCD monitor (Samsung Electronics, Yeongtong, Suwon, South Korea) at a refresh rate of 60 Hz. Stimulus presentation was synchronized with imaging using transistor-transistor logic signal of image acquisition timing and its counter board (USB-6501, National Instruments, Austin, TX, USA).

First, the retinotopic position was determined using moving grating patches (contrast: 99.9%, spatial frequency: 0.04 cycles/degree, temporal frequency: 2 Hz). We first determined the coarse retinotopic position by presenting a grating patch with a 50-degree diameter at each 5 ×3 position covering the entire monitor. Then, a grating patch with a 20-degree diameter was presented at each 4 ×4 position covering an 80 ×80-degree space to fine-tune the position. The retinotopic position was defined as the position with the highest response.

Natural images (200, 1200, or 2200 images, 512 ×512 pixels) were obtained from the van Hateren Database [54] and McGill Calibrated Colour Image Database [55]. After each image was gray-scaled, it was preprocessed such that its contrast was 99.9% and its mean intensity across pixels was at an intensity level of approximately 50%, and then masked with a circle with a 60-degree diameter. The stimulus presentation protocol consisted of 3-12 sessions. In one session, images were ordered pseudo-randomly, and each image was flashed three times in a row. Each flash was presented for 200 ms with 200-ms intervals between flashes in which a gray screen was presented.

### Acquisition of simulated data

The following types of artificial cells were simulated in this study: simple, complex, and rotation-invariant cells. A simple cell was modeled using a “linear-nonlinear” cascade formulated as shown below where the response to a stimulus was defined as the dot product between the stimulus image *s* and a Gabor-shaped filter *f*_*1*_, followed by a rectifying nonlinearity [2] and a Gaussian noise (Fig 2A).

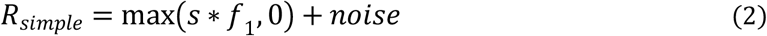

A complex cell was modeled using an energy model with two subunits [36,37]. In this model, each subunit calculated the dot product between the stimulus image s and a Gabor-shaped filter *f*_*1*_, *f*_*2*_. Then, the outputs of these two subunits were squared, summed together, and the squared root was taken. Finally, a Gaussian noise was added to define the response (Fig 2B). Here, the Gabor-shaped filters used in this model had identical amplitude, position, size, spatial frequency, and orientation; the phase was shifted by 90 degrees. Note that this procedure, formulated as follows, can also be viewed as a “linear-nonlinear-linear-nonlinear” cascade [30,56].

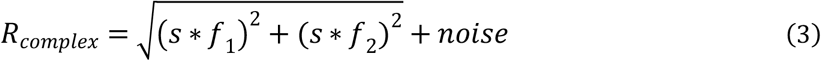

A rotation-invariant cell was modeled using 36 subunits. The *i*-th subunit (1 < *i* < 36) calculated the dot product between the stimulus image s and a Gabor-shaped filter *f*_*i*_. After the maximum of the outputs of the subunits was taken, a Gaussian noise was added to define the response (Fig 3A). Here, the Gabor-shaped filters used in this model *f*_*i*_ had identical amplitude, position, size, spatial frequency, and phase; the orientation of the *i*-th subunit was 5 (*i* - 1) degree.

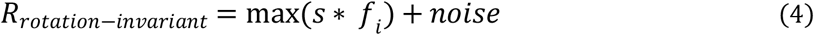

We simulated 30 simple cells, 70 complex cells, and 10 rotation-invariant cells. For each cell simulation, we performed 4 trials with a different random noise. The stimuli used in these three models were identical to the stimuli used in the acquisition of real neural data (2200 images), which were down-sampled to 10 ×10 pixels. The Gabor-shaped filter used in these models, a product of a two-dimensional Gaussian envelope and a sinusoidal wave, was formulated as follows:

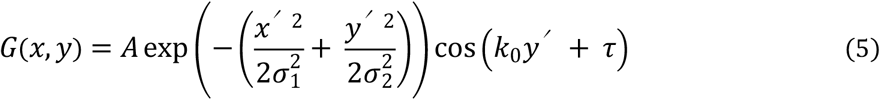

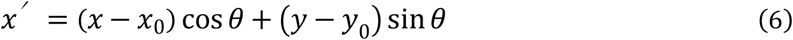

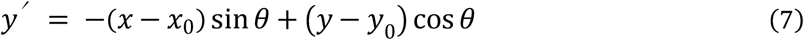

where *A* is the amplitude, *σ*_*1*_ and *σ*_*2*_ are the standard deviations of the envelopes, *k*_*0*_ is the frequency, *τ* is the phase, (*x*_*0*_, *y*_*0*_) is the center coordinate, and *θ* is the orientation. The parameters for *f*_*1*_ of simple cells and complex cells were sampled from a uniform distribution over the following range: *0.1* ≤ *x*_*0*_ / *L*_*x*_ ≤ *0.9*, *0.1* ≤ *y*_*0*_ / *L*_*y*_ ≤ *0.9*, *0* ≤ *A* ≤ *1*, *0.1* ≤ *σ*_*1*_ / *L*_*x*_ ≤ *0.2*, *0.1* ≤ *σ*_*2*_ / *L*_*y*_ ≤ *0.2*, *π/3* ≤ *k*_*0*_ ≤ *π*, *0* ≤ *θ* ≤ *2π*, and *0* ≤ *τ* ≤ *2π*, where *L*_*x*_ and *L*_*y*_ are the size of the stimulus image in the x and y dimension, respectively. The parameters for *f*_*1*_ of rotation-invariant cells were sampled from a uniform distribution over the following range: *0* ≤ *A* ≤ *1*, *0.15* ≤ *σ*_*1*_ / *L*_*x*_ ≤ *0.2*, *0.15* ≤ *σ*_*2*_ / *L*_*y*_ ≤ *0.2*, *π/3* ≤ *k*_*0*_ ≤ *2/3π* and *0* ≤ *τ* ≤ *2π*. *x*_*0*_, *y*_*0*_ and *θ* were set as *L*_*x*_/2, *L*_*y*_/2, and 0, respectively.

The noise was randomly sampled from a Gaussian distribution with a mean of zero and standard deviation of one, which resulted in trial-to-trial variability similar to that of real data.

### Data preprocessing

Data analyses were performed using Matlab (Mathworks, Natick, MA, USA) and Python (2.7.13, 3.5.2, and 3.6.1). For real neural data, images were phase-corrected and aligned between frames [57]. To determine regions of interest (ROIs) for individual cells, images were averaged across frames, and slow spatial frequency components were removed from the frame-averaged image with a two-dimensional Gaussian filter whose standard deviation was approximately five times the diameter of the soma. ROIs were first automatically identified by template matching using a two-dimensional difference-of-Gaussian template and then corrected manually. SR101-positive cells, which were considered putative astrocytes [58], were removed from further analyses. The time course of the fluorescent signal of each cell was calculated by averaging the pixel intensities within an ROI. Out-of-focus fluorescence contamination was removed using a method described previously [59,60]. The neuronal response to each natural image was computed as the difference between averaged signals during the last 200 ms of presentation and averaged signals during the interval preceding the image presentation.

For both real data and simulated data, responses were averaged across all trials and scaled such that the values were between zero and one. Natural images used in further analyses were down-sampled to 10 ×10 pixels. We finally standardized the distribution of each pixel by subtracting the mean and then dividing it by the standard deviation.

### Encoding models

Encoding models were developed for each cell. An L1-regularized linear regression model (Lasso), L2-regularized linear regression model (Ridge), and SVR with radius basis function kernel were implemented using the Scikit-learn (0.18.1) framework [61]. The hyperparameters of these encoding models were optimized by exhaustive grid search with 5-fold cross-validation for data of 10 real V1 neurons. The optimized hyperparameters were as follows: the regularization coefficients of Lasso and Ridge were 0.01 and 10^4^, respectively, and the kernel coefficient and penalty parameter of SVR were both 0.01. The HSM was implemented as previously proposed [31] with hyperparameters identical to the ones used in the study.

CNNs were implemented using the Keras (2.0.3 and 2.0.6) and Tensorflow (1.1.0 and 1.2.1) framework [62]. A CNN consisted of the input layer, several hidden layers (convolutional layer, pooling layer, or fully connected layer), and the output layer. The activation of a convolutional layer was defined as the rectified linear (ReLU) [63] transformation of a two-dimensional convolution of the previous layer activation. Here, the number of convolutional filters in one layer was 32, the size of each filter was (3, 3), the stride size was (1, 1), and valid padding was used. The activation of a pooling layer was 2×2 max-pooling of the previous layer activation, and valid padding was also used. The activation of a fully connected layer was defined as the ReLU transformation of the weighted sum of the previous layer activation. If the previous layer had a two-dimensional shape, the activation was flattened to one dimension. The activation of the output layer was the sigmoidal transformation of the weighted sum of the previous layer. The size of the mini batch, dropout [64] rate, type of optimizer (stochastic gradient descent (SGD) or Adam [65]), learning rate decay coefficient of SGD, and number and types of hidden layers (convolutional, max-pooling, or fully connected) were optimized with 5-fold cross-validation for the data of 10 real V1 neurons. The optimized hyperparameters of CNN were as follows: the size of the mini batch was 5 or 30 (depending on the size of the dataset), the dropout rate of fully connected layers was 0.5, the optimizer was SGD, the learning rate decay coefficient was 5 × 10^−5^, and the hidden layer structure was 4 successive convolutional layers and one pooling layer, followed by one fully connected layer (Fig 1). Other hyperparameters were fixed.

The training was formulated as follows:

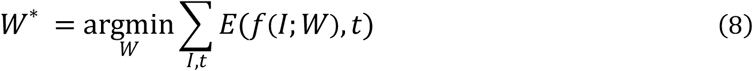

where *I* is an image, *t* is the response, *W* is the parameters, and *f* is the model. *E* is the loss function defined as the mean squared error between the predicted responses and actual responses in the training dataset. The prediction accuracy was defined as the Pearson correlation coefficient between the predicted responses and actual responses. The training procedures of CNNs were as follows. First, the training data were subdivided into data used to update the parameters (90% of training data) and data used to monitor generalization performances (10% of training data: validation set). After the parameters were initialized by sampling from Glorot uniform distributions [66], they were updated iteratively by backpropagation [67], which was performed to minimize the loss function in either a SGD or Adam manner. SGD was formulated as follows:

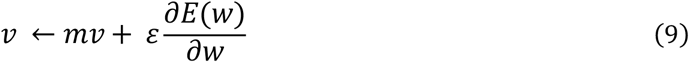

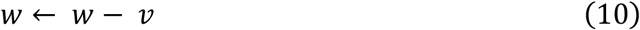

where *w* is the parameter we want to update, *m* is the momentum coefficient (0.9), *v* is the momentum variable, *ε* is the learning rate (initial learning rate was 0.1), and *E(w)* is the loss with respect to the batched data. Adam was formulated as previously suggested [65]. The training iterations were stopped upon saturation of the prediction accuracy for the validation set.

The response prediction accuracy of each encoding model was evaluated in a 5-fold cross-validation manner for each cell not used for hyperparameter optimizations. To quantify the nonlinearity of each cell, we defined a nonlinearity index for each cell by comparing the response prediction accuracy of Lasso and CNN in the following way:

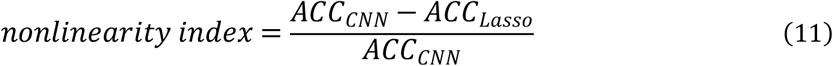

where *ACC*_*CNN*_ and *ACC*_*Lasso*_ are the response prediction accuracy of CNN and Lasso, respectively.

### RF estimation

Nonlinear RFs were estimated from trained CNNs using a regularized version of a maximization-of-activation approach [23,24]. Cells with a CNN prediction accuracy ≤ 0.3 were omitted from this analysis. First, CNN was trained using all data for each cell. Then, starting with a randomly initialized image, an image *I* was updated iteratively (10 times) by gradient ascent to maximize the following objective function *E(I)*:

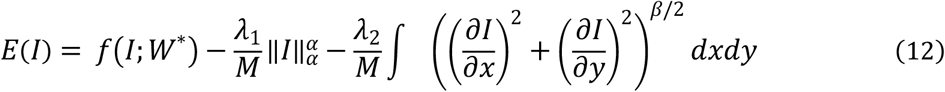

where *f* is the trained CNN model; *W*\* is the trained parameters, which is fixed in this procedure; *λ*_*1*_, *λ*_*2*_, *α*, and *β* are the regularization parameters, which are fixed as 10, 2, 6, and 1, respectively; and *M* is the size of the image. The second and third terms are regularization terms to minimize the α-norm and total variation [26] of the image, respectively. The RMSprop algorithm [68] was used as the gradient ascent formulated as follows:

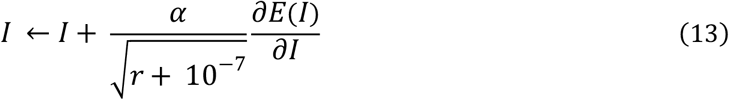

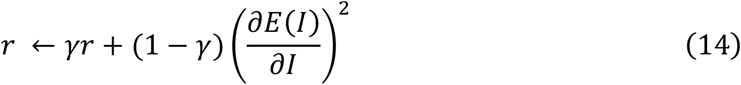

where *γ* is the decay coefficient (0.95) and *α* is the learning rate (1.0). The generated image was finally processed such that its mean was zero and standard deviation was one (RF image). To confirm that the generated RF image maximally activates the output layer, the whole process was repeated independently until we generated an image to which the predicted response was high (for most cells, > 95% of the maximum response of the actual data of each cell). Note that for representative cells (Figs 2D, 2E, 3C, and 4B), the predicted responses to the generated RF images were > 99% of the maximum response of the actual data.

To quantitatively assess the generated RF images, we fitted each RF image with a Gabor kernel *G(x, y)* using sequential least-squares programming implemented in Scipy (0.19.0). A Gabor kernel, a product of a two-dimensional Gaussian envelope and a sinusoidal wave, was formulated as follows:

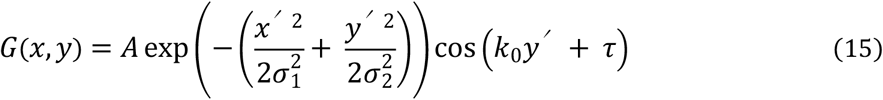

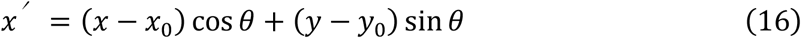

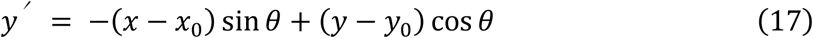

where *A* is the amplitude, *σ*_*1*_ and *σ*_*2*_ are the standard deviations of the envelopes, *k*_*0*_ is the frequency, t is the phase, (*x*_*0*_, *y*_*0*_) is the center coordinate, and *θ* is the orientation. The goal of fitting was to minimize the pixelwise absolute error between the RF image and a Gabor kernel. This optimization was started with seven different initial *x*_*0*_ and seven different initial *y*_*0*_ to ensure that the optimization fell in the global minima. In addition, to create a reasonable Gabor kernel, we set bounds for some of the parameters: *0* ≤ *x*_*0*_ / *L*_*x*_ ≤ *1*, *0* ≤ *y*_*0*_ / *L*_*y*_ ≤ *1*, *0* ≤ *0* ≤ *σ*_*1*_ / *L*_*x*_ ≤ *0.2*, *0* ≤ *σ*_*2*_ / *L*_*y*_ ≤ *0.2*, and *π/3* ≤ *k*_*0*_ *τ*, where *L*_*x*_ and *L*_*y*_ are the size of the RF image in the x and y dimension, respectively. The accuracy of Gabor fitting was evaluated by the pixelwise Pearson correlation coefficient between the original RF image and the fitted Gabor kernel.

Linear RF images were created by a regularized pseudoinverse method described previously [38]. The regularization parameter was optimized for each cell by exhaustive grid search in a 10-fold cross-validation manner. For each value in the grid, responses to the held-out test data were predicted using the created RF image. Prediction accuracy was calculated as the Pearson correlation coefficient between the predicted responses and actual responses. The linear RF image was created using the value with the highest prediction accuracy as the regularization parameter.

### Quantification of shift-invariance (complexness)

To distinguish between simple cells and complex-like cells, we then created a “shifted image set”, which contained CNN RF images that were shifted with respect to one another, selected from the 100 CNN RF images. For this purpose, a zero-mean normalized cross correlation (ZNCC) was calculated for every pair of RF images (*I*_*1*_, *I*_*2*_):

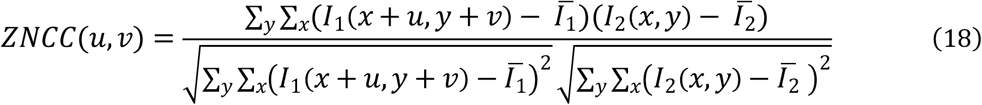

where (*u*, *v*) is a pixel shift and 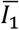 is the mean of *I*_*1*_. If the ZNCC was above 0.95 for a (*u*, *v*) pair ((*u*, *v*) ≠ (0, 0)), these two RF images were defined as shifted to each other by (*u*, *v*) pixels. Then, for each pair of shifted RF images, we calculated the shift distance as the maximum length of (*u*, *v*) vectors projected orthogonally to the Gabor orientation. Finally, starting with the two RF images with the largest shift distance, we iteratively collected RF images that were shifted from the already collected RF images to create the “shifted image set”. If none of the 100 RF images were shifted to another, the “shifted image set” consisted of the RF image with the highest predicted response.

A simple model and complex model were created for each cell as follows (Fig 7). In the simple model, the response to a stimulus image was predicted as the normalized dot product between the stimulus image and one RF image selected from the “shifted image set”. The RF image that yielded the best prediction accuracy was chosen and used for all stimulus images. In the complex model, the response to a single stimulus image was predicted as the maximum of the normalized dot products between the stimulus image and RF images in the “shifted image set”. The RF image with the maximal dot product was selected for each stimulus image separately. The prediction accuracy for each model was quantified as the Pearson correlation coefficient between the predicted responses and actual responses among all stimulus-response datasets. Finally, the complexness index for each cell was defined by

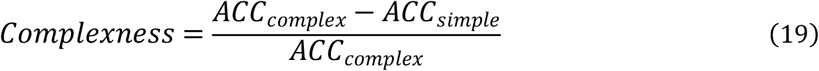

where *ACC*_*simple*_ and *ACC*_*complex*_ are the response prediction accuracy of the simple model and complex model, respectively. Cells with the Gabor fitting accuracy ≤ 0.6, *ACC*_*simple*_ < 0, or *ACC*_*complex*_ < 0 were omitted from this analysis.

### Spatial organizations of simple cells and complex-like cells

The spatial organizations of simple cells and complex-like cells were evaluated in two ways. First, for each pair of neurons, we calculated the in-between cortical distance and the difference in complexness. A relationship between the cortical distances and the complexness differences is indicative of a spatial organization [57]. Second, we calculated the cumulative distributions of the in-between cortical distances for all pairs of simple cells and for all pairs of complex-like cells. To statistically evaluate the cumulative distributions, we permuted the cell positions 1000 times independently for each plane. For each permutation, cell positions of simple cells were randomly sampled from original cell positions of simple and complex-like cells. Other positions were allocated for complex-like cells. After the cell positions were determined, the cumulative distributions of the in-between cortical distances were calculated. After repeating this procedure independently 1000 times for each plane, the first and 99^th^ percentiles of the permuted cumulative distributions were calculated for the significance levels.

## Acknowledgements

We thank all members of the Ohki laboratory, especially Ms. T. Inoue, Y. Sono, A. Ohmori, A. Honda, and M. Nakamichi for animal care. This work was supported by grants from Brain Mapping by Integrated Neurotechnologies for Disease Studies (Brain/MINDS), Japan Agency for Medical Research and Development (AMED) (to K.O.); International Research Center for Neurointelligence (WPI-IRCN), Japan Society for the Promotion of Sciences (JSPS) (to K.O.); Core Research for Evolutionary Science and Technology (CREST), AMED (to K.O.); Strategic International Research Cooperative Program (SICP), AMED (to K.O.); JSPS KAKENHI (grant number 25221001 and 25117004 to K.O. and 15K16573 and 17K13276 to T.Y.); the Ichiro Kanehara Foundation for the Promotion of Medical Sciences and Medical Care (to T.Y.); and the Uehara Memorial Foundation (to T.Y.). J.U. was supported by the Takeda Science Foundation and Masayoshi Son Foundation.

## Author contributions

J.U., T.Y., and K.O. designed the study; T.Y. performed the experiments; J.U., T.Y., and K.O. analyzed the data; and J.U., T.Y., and K.O. wrote the paper.

